# Structural and Functional analysis of the GABARAP Interaction Motif (GIM)

**DOI:** 10.1101/083659

**Authors:** Vladimir V. Rogov, Alexandra Stolz, Arvind C. Ravicahandran, Alexander Law, Hironori Suzuki, Andreas Kniss, Diana O. Rios-Szwed, Frank Löhr, Soichi Wakatsuki, Volker Dötsch, Ivan Dikic, Renwick C.J. Dobson, David G. McEwan

## Abstract

Through the canonical LC3 interaction motif (LIR), [W/F/Y]-X_1_-X_2_-[I/L/V], protein complexes are recruited to autophagosomes to perform their functions as either autophagy adaptors or receptors. How these adaptors/receptors selectively interact with either LC3 or GABARAP families remains unclear. Herein, we determine the range of selectivity of 30 known core LIR motifs towards LC3s and GABARAPs. From these, we define a <u>**G**</u>ABARAP <u>**I**</u>nteraction <u>**M**</u>otif (GIM) sequence (W/F-V-X_2_-V) that the adaptor protein PLEKHM1 tightly conforms to. Using biophysical and structural approaches, we show that the PLEKHM1-LIR is indeed eleven-fold more specific for GABARAP than LC3B. Selective mutation of the X_1_ and X_2_ positions either completely abolished the interaction with all LC3 and GABARAPs or increased PLEKHM1-GIM selectivity 20-fold towards LC3B. Finally, we show that conversion of the canonical p62/SQSTM1-LIR into our newly defined GIM, by introducing two valine residues, enhances p62/SQSTM1 interaction with endogenous GABARAP over LC3B. The identification of a GABARAP-specific interaction motif will aid the identification and characterization of the continually expanding array of autophagy receptor and adaptor proteins and their *in vivo* functions.

## Introduction

Autophagy is an alternative catabolic process that works alongside the proteasome for the degradation of cellular material. Such cargo can include protein aggregates, damaged organelles, intracellular pathogens, metabolic substrates and ferritin aggregates [1-4]. At the heart of the autophagy pathway, are ubiquitin-like proteins that, despite sharing little primary sequence with ubiquitin, contain an ubiquitin-like fold [5]. The most characterized of these ubiquitin-like modifiers is the *Saccharomyces cerevisiae* Atg8 protein. Unlike *Saccharomyces cerevisiae*, however, there are six Atg8 homologues in mammals (mammalian Atg8s; mATG8s) that, presumably, have distinct or overlapping functions: MAP1LC3A (microtubule-associated protein light chain 3 alpha; LC3A), LC3B, LC3C, GABARAP (γ-aminobutyric acid receptor-associated protein), GABARAP-L1 and GABARAP-L2/GATE-16 [6].

All six mATG8s are essential for autophagy, are conjugated to autophagosomes and serve to recruit two broad classes of molecules: autophagy receptors and autophagy adaptors. Autophagy receptors interact directly with mATG8s on the inner autophagosomal membrane and provide a vital link between the autophagosomal isolation membrane and cargo to be sequestered and delivered to the lysosome for degradation, for example, protein aggregates (p62 [7]; NBR1 [8]; Cue5 [9]) or intracellular pathogens (OPTN [10]; NDP52 [11]; TAX1BP1 [12]). Additionally, organelles such as ER (FAM134B [4]), mitochondria (Nix/BNIP3L [13]) as well as ferritin (NCOA4 [14]) can be specifically targeted by autophagy receptors. On the cytosolic facing membrane, mATG8s interact with adaptor proteins that regulate autophagosome formation (ULK1/2 [15]), autophagosome transport (FYCO1 [16]), cross talk with the endocytic network (TBC1D5 [17]) and autophagosome fusion with the lysosome (PLEKHM1 [18]), but are not themselves degraded by autophagy. Autophagy ubiquitin like modifiers can also act as signalling scaffolds to attract diverse complexes, such as GABARAP-mediated recruitment of CUL3-KBTBD6/KBTBD7 ubiquitin ligase complex to a membrane-localized substrate, TIAM1 [19]. One essential common feature of all adaptors and receptors is the presence of a LC3 interaction region (LIR; also known as LC3 interaction motif (LIM) or Atg8 interaction motif (AIM)).

With some known exceptions (“atypical LIRs/LIMs”), such as NDP52 [11], TAX1BP1 [20] and the dual LIR/UFIM (UFM1-Interaction Motif) in UBA5 [21], the majority of LIRs contain a core Θ-X_1_-X_2_-Γ motif, where Θ is an aromatic residue (W/F/Y), and Γ is a large hydrophobic residue (L/V/I). Structural studies have shown that the side-chains of the aromatic residue (Θ) within the core LIR motif are placed deep inside of a hydrophobic pocket (HP1) on the Atg8/LC3/GABARAP surface, formed between α-helix 2 and β-strand 2, while side-chains of the hydrophobic LIR residues (Γ) occupies a second hydrophobic pocket (HP2) between β-strand 2 and α-helix 3 (reviewed in [3, 22, 23]). Acidic and phosphorylatable serine/threonine residues n-terminal, and occasionally c-terminal, of the core LIR/AIM can contribute to the stabilization of LIR-mATG8 interactions [24-26]. There is growing evidence that the function of the autophagy adaptors and receptors are closely linked to their interaction with specific LC3/GABARAP family members and their distinct role in the pathway [18, 27, 28]. The presence of six similar LC3/GABARAP proteins also points towards their specific functions within the pathway; for example, at the formation and closure of the nascent phagophore during autophagosome formation [28]. Therefore, despite having similar sequences, there is a clear selectivity and divergence of function between the six mATG8s. However, as yet, there has been no identification of an LC3- or GABARAP-subfamily selective LIR motif.

In order to address the issue of selectivity, we implemented a peptide based assay to screen 30 validated LIR sequences against all LC3 and GABARAP proteins, with the main focus on positions X_1_ and X_2_ located within the core Θ-X_1_X_2_-Γ sequence. We identified 13 GABARAP-preferring LIR sequences, and analysed the PLEKHM1-LIR in detail to understand the driving forces of the observed specificity. We propose that residues within the classical LIR sequence, particularly at the X_1_ and X_2_ positions, help to define subfamily selectivity and that we can alter selectivity by changing residues in these positions. These data will help define the interaction motifs as either AIM (Atg8), LIR (LC3) or GIM (GABARAP) and develop our understanding of subfamily specific interactions and their functional consequences.

## Results

### LIR motifs of known autophagy receptors and adaptors feature mATG8 specificity

A high number of autophagy receptors or adaptor structures have been reported, yet the basis for their selective interaction with individual members of the ATG8 family is not well understood. We speculated whether the LIR motif alone is able to confer selectivity towards a mATG8 subfamily and if we could derive a subfamily motif from analysis of known mATG8 interaction partners. To address this question, we screened an array of peptides (presented in **Expanded View Figure EV1A** and described in Material and Methods) with the LIR sequences of 30 known and validated autophagy receptors and adaptors (Table 1) against all six human mATG8s for binding (Figure 1A and **Expanded View Figures EV1B-C**). In brief, biotinylated peptides were immobilized on streptavidin coated 96-well plates and incubated with His_6_-tagged mATG8 proteins. After washing steps, peptide bound mATG8 was detected in an ELISA reader using anti-His antibodies directly conjugated to HRP (horse radish peroxidase) (**Expanded View Figure EV1A**).

**Figure 1:**
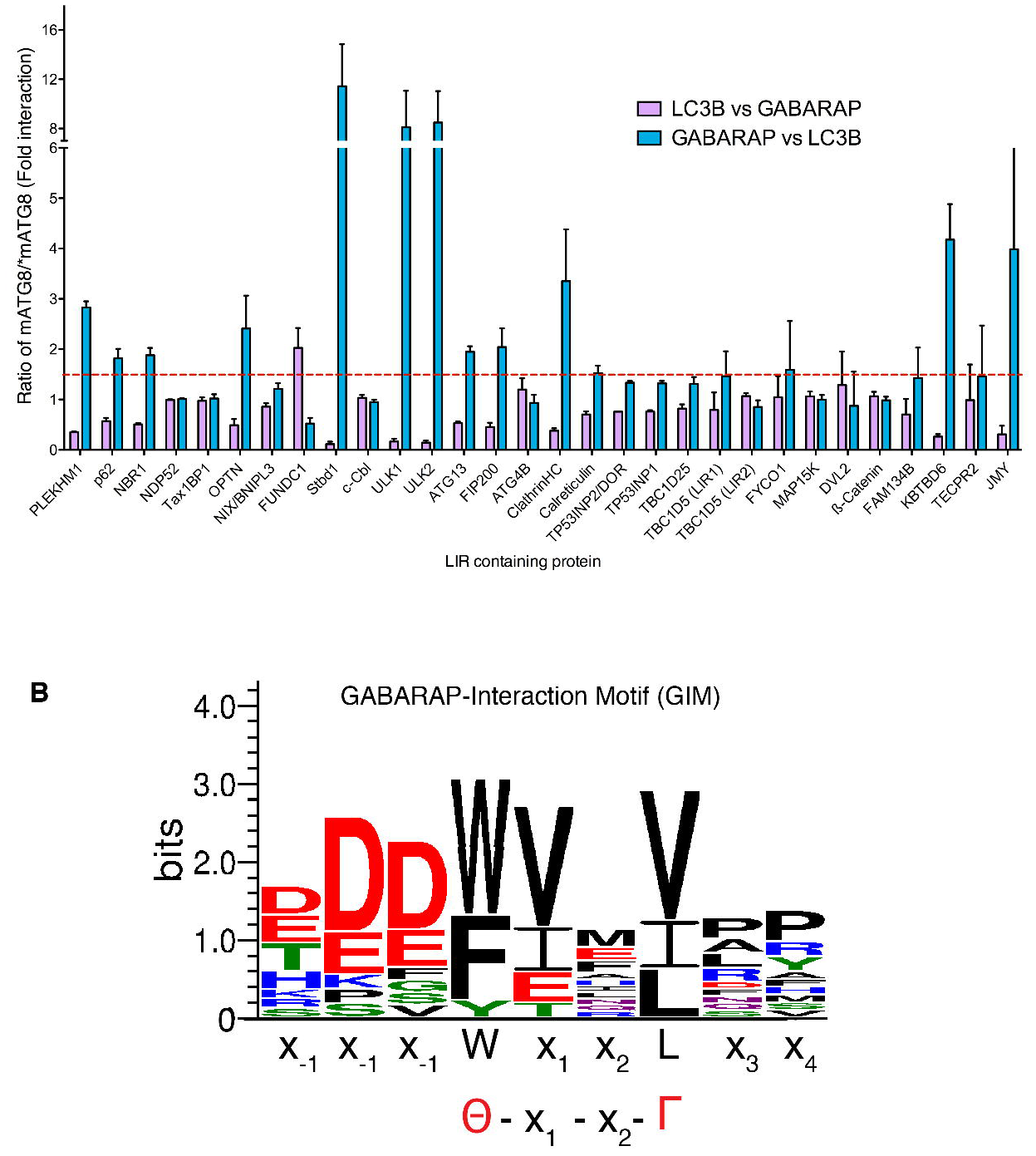
Defining a mATG8-subfamily specific interaction motif. **A.** Interaction profile of 30 biotinylated LIR peptides from various proteins against 6xHis-tagged-LC3B and 6xHis-tagged GABARAP. Results for each LIR interactions were expressed as either absorbance of LC3B divided by absorbance of GABARAP (purple bars) or absorbance of GABARAP divided by absorbance of LC3B (blue bars) to define whether each LIR shows preference towards either LC3- or GABARAP-family proteins. Dashed red line depicts 1.5 fold change cut-off. Values are mean of n=3 independent experiments ± S.E.M. **B.** WebLogo generated from 14 sequences that showed preference towards GABRAP versus LC3B interaction.

**Table 1.**
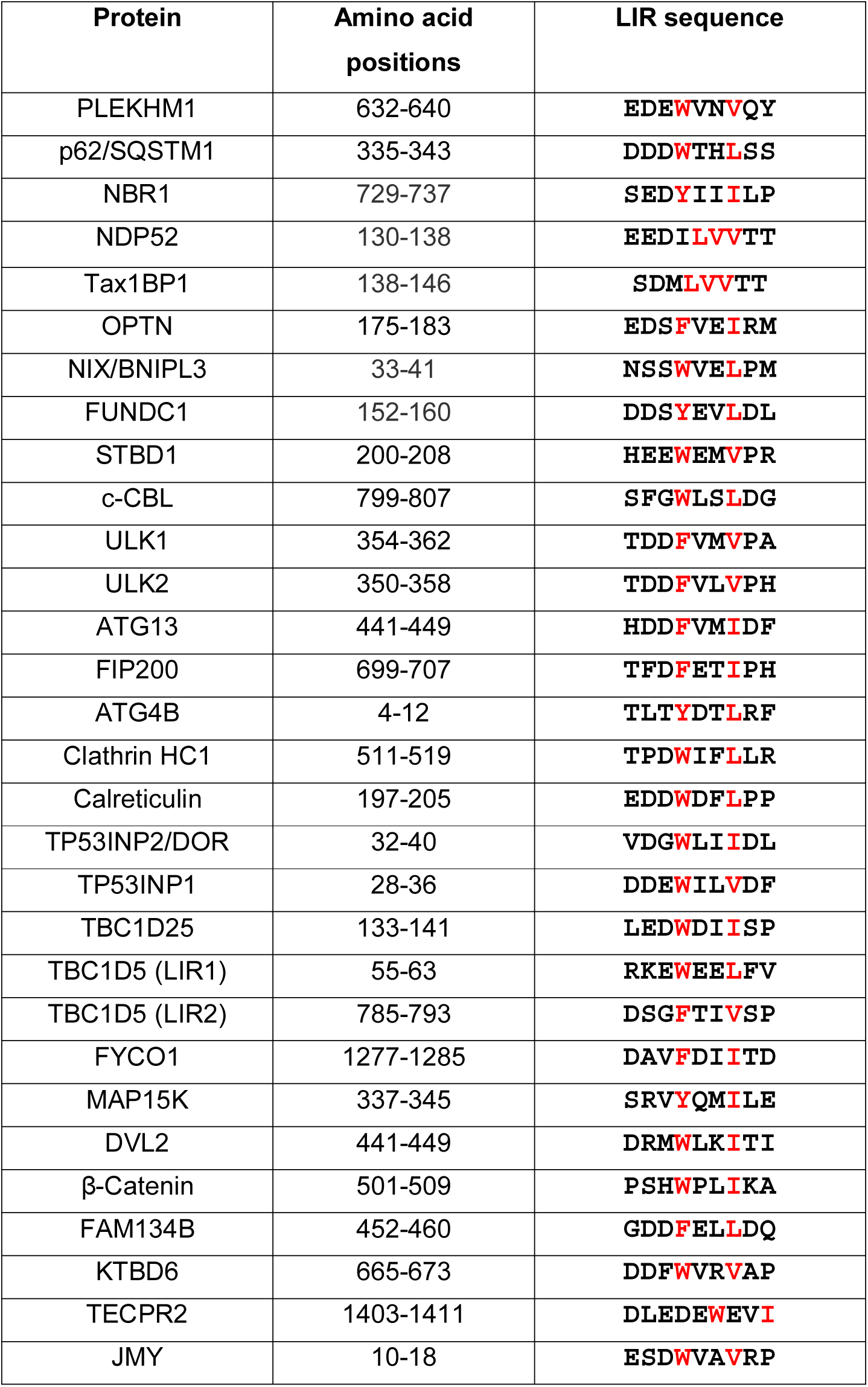
LIR sequences of known mATG8 proteins tested for interactions with all LC3 and GABARAP proteins.

Due to the wide range of affinities of various LIR sequences towards the LC3/GABARAP proteins, we have normalized our results by dividing values for LC3B interaction by the corresponding value for interaction with GABARAP (Figure 1A, purple bars) and vice versa (Figure 1A, blue bars) to highlight the potential subfamily selectivity of each LIR sequence tested. We classified ratios greater than 1.5 fold as an indication of a preferential interaction towards that particular LC3 or GABARAP family member. Out of the 30 LIRs tested, 12 (40%) showed selectivity towards GABARAP over LC3 subfamily (Table 2) and only one LIR, FUNDC1, preferentially interacted with the LC3 group (Figure 1A). These results are consistent with previously published data, with for example, ULK1/ULK2 and KBTDB6, showing a clear specificity towards GABARAP versus LC3B [19, 24]. Using this information, we generated a sequence plot (Figure 1B) to ascertain whether there were any common sequence features of the GABARAP-specific interaction proteins. In addition to the 12 sequences identified in this experiment as preferential GABARAP-sub family interactors, we also included known GABARAP interactors that were not included in our screen (ALFY and KBTBD7). We found that the fourteen LIR sequences had a high frequency of valine in the X_1_ position (8 out of 14, 57%) with another 3 (21%) having an isoleucine (Table 2), indicating that both V and I at position X_1_ may represent a distinguishing feature of GABARAP selective LIR sequences. The previously identified PLEKHM1-LIR [18] has a high degree of similarity to this sequence. PLEKHM1 can interact with all LC3s in a GST-pull down assay [18], but we detected a clear preference for binding to GABARAP and GABARAP-L2 over LC3B and LC3C (Figure 1A and **Expanded View Figure EV1B-C**). This result supports a function of PLEKHM1 as an adaptor and not a receptor protein.

**Table 2.**
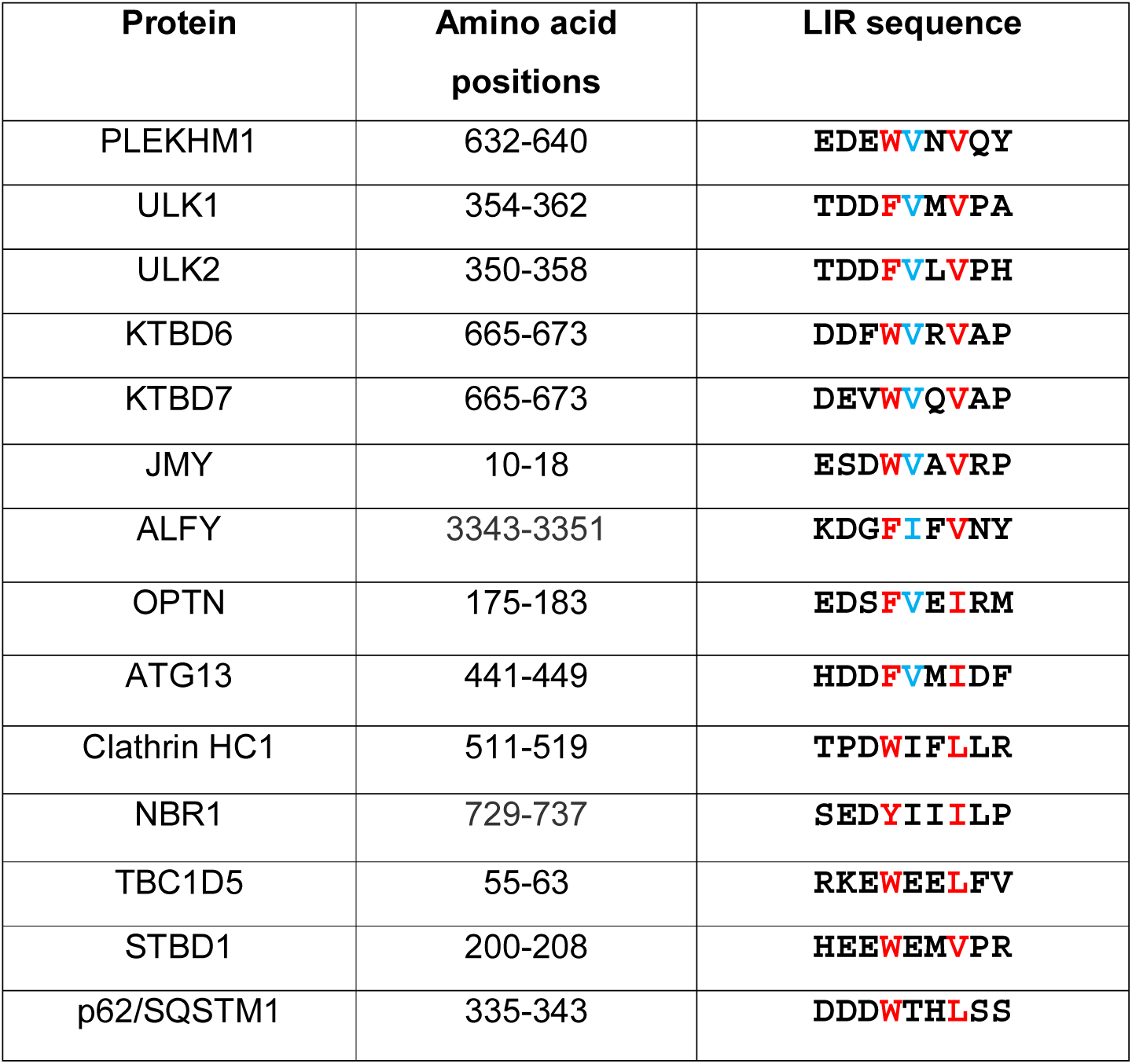
LIR sequences of 14 GABARAP-selective interacting proteins.

### PLEKHM1 interacts preferentially with GABARAP-family proteins

To further characterize the LIR sequences with preferential binding to GABARAP-subfamily proteins, we employed biochemical and biophysical techniques to study interactions of the PLEKHM1-LIR with all six mammalian LC3/GABARAP proteins.

Isothermal titration calorimetry (ITC) experiments titrating purified PLEKHM1-LIR peptide to all six mATG8s (LC3A, LC3B, LC3C, GABARAP, GABARAP-L1 and GABARAP-L2) revealed K_D_ values in the μM range (Figure 2A and Table 3). Consistent with the previous data (Figure 1A), the GABARAP-family proteins had significantly lower K_D_ values compared to the LC3 family. Indeed, the K_D_ of GABARAP (0.55 μM) with the PLEKHM1-LIR peptide is approximately 8 times lower compared to LC3A (4.22 μM) and approximately 11 times lower compared to LC3B (6.33 μM) (Figure 2A and **Expanded View Figure EV2A**). In addition, we performed NMR experiments titrating ^15^N-labelled LC3A, LC3B, GABARAP-L1 and GABARAP-L2 samples (as representative members of LC3- and GABARAP-subfamilies) with the PLEKHM1-LIR peptide. In agreement with the ITC data we observed slow exchange behavior of resonance of the GABARAP subfamily proteins and intermediate exchange for the LC3 subfamily proteins upon titration (**Expanded View Figure EV2B**). We mapped the chemical shift perturbations (CSP) on the (**Expanded View Figure EV2C**) structures (**Expanded View Figure EV2D**) of all four proteins used in this experiment, revealing a high degree of similarity in the CSP patterns. Most affected are the backbone HN resonances of residues forming the hydrophobic pockets 1 and 2 (HP1 and HP2, highlighted in **Expanded View Figure EV2D**), and β-strand 2 which participates in formation of the intermolecular β-sheet between mATG8 proteins and LIR sequences [19, 22, 23, 29, 30].

**Figure 2.**
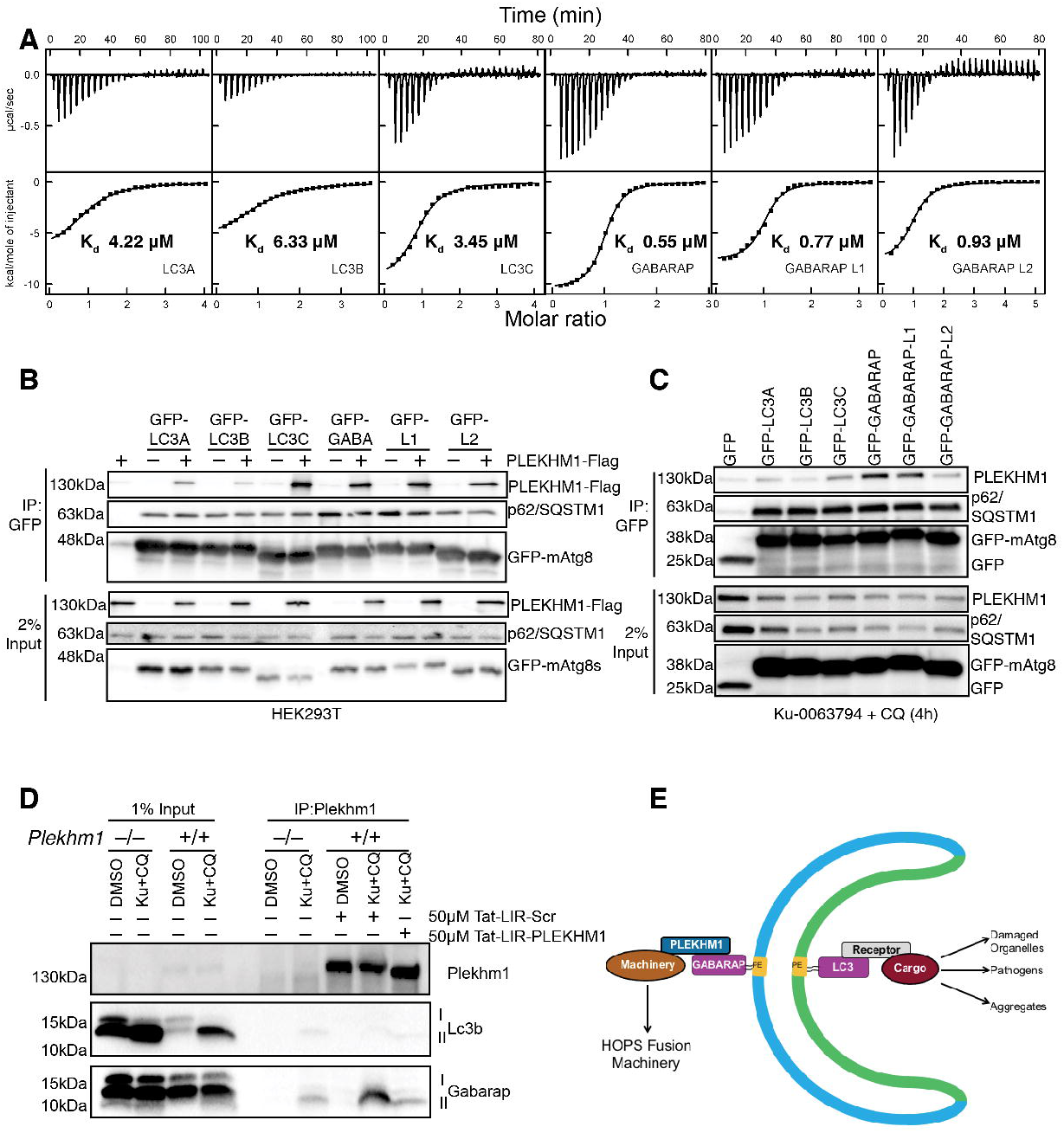
PLEKHM1 preferentially interacts with GABARAP *in vitro* and *in vivo*. **A.** ITC titrations of PLEKHM1-LIR peptide into LC3-family proteins (top panel) and GABARAP-family proteins (bottom panel). The top diagrams in each ITC plot display the raw measurements and the bottom diagrams show the integrated heat per titration step. Best fit is presented as a solid line. **B.** GFP-tagged LC3/GABARP proteins were expressed alone or with PLEKHM1-WT-Flag in HEK293T cells and immunoprecipitated using GFP-Trap beads and blotted for the presence or absence of PLEKHM1 (Anti-Flag tag). **C.** GFP-LC3/GABARAPs were overexpressed in HeLa cells and treated for 4 h with KU-0063794 (10μM) plus chloroquine (20μM), immunoprecipitated with GFP-Trap beads and blotted for the presence of endogenous PLEKHM1. **D.** *Plekhm1*^+/+^ *or Plekhm1*^−/−^ mouse embryonic fibroblasts were either treated with vehicle (DMSO) or treated for 4 h with KU-0063794 (10μM) plus chloroquine (20μM). Samples were lysed in NP-40 lysis buffer endogenous PLEKHM1was immunoprecipitated in the presence of 50μM Tat-PLEKHM1-LIR peptide (Tat-KVRPQQ<u>**EDEWVNV**</u>QYPDQPE) or 50μM Tat-Scrambled (Scr) PLEKHM1-LIR peptide (Tat-VQEQQEPPPVKNYDVEQWDR). Samples were then immunoblotted for the presence of endogenous PLEKHM1, LC3B and GABARAP proteins. **E.** Schematic representation of PLEKHM1 interaction on Autophagosomes with GABARAP on cytosolic facing membrane.

**Table 3:** Thermodynamic parameters of interactions between LC3/GABARAP proteins and PLEKHM1-LIR peptide.

| | $\Delta H$<br>kcal mol<sup>-1</sup> | $\Delta S$<br>cal mol<sup>-1</sup> K<sup>-1</sup> | $-T\Delta S$<br>kcal mol<sup>-1</sup> | $\Delta G$<br>kcal mol<sup>-1</sup> | $K_A$<br>*10<sup>6</sup> M<sup>-1</sup> | $K_D$<br>μM | **N** |
| --- | --- | --- | --- | --- | --- | --- | --- |
| PLEKHM1-LIR WT |  |  |  |  |  |  |  |
| LC3A | -7±0.2 | +1.17 | -0.35 | -7.33 | 0.24±0.02 | 4.22 | 0.98±0.02 |
| LC3B | -5.8±0.2 | +4.27 | -1.27 | -7.09 | 0.16±0.01 | 6.33 | 1.06±0.01 |
| LC3C | -8.3±0.2 | -2.83 | +0.84 | -7.48 | 0.29±0.02 | 3.45 | 0.99±0.02 |
| G<sub>ABARAP</sub> | -10.6±0.1 | -6.92 | +2.06 | -8.54 | 1.8±0.1 | 0.55 | 1.00±0.01 |
| G<sub>ABARAP</sub>-L1 | -7.8±0.1 | +1.94 | -0.58 | -8.35 | 1.3±0.1 | 0.77 | 1.00±0.01 |
| G<sub>ABARAP</sub>-L2 | -6.1±0.1 | +7.00 | -2.09 | -8.23 | 1.07±0.03 | 0.93 | 1.05±0.01 |
| PLEKHM1-LIR W635A |  |  |  |  |  |  |  |
| LC3B | -0.8±0.1 | +15.4 | -4.59 | -5.44 | 0.01±0.01 | >100 | 1* |
| G<sub>ABARAP</sub> | -1.1±0.1 | +14.4 | -4.29 | -5.40 | 0.01±0.01 | >100 | 1* |
| PLEKHM1-LIR V636G |  |  |  |  |  |  |  |
| LC3B | -1.7±0.1 | +12.4 | -3.70 | -5.42 | 0.01±0.01 | >100 | 1* |
| G<sub>ABARAP</sub> | -7.5±0.1 | -5.66 | +1.69 | -5.83 | 0.02±0.00 | 52 | 1* |
| PLEKHM1-LIR N637G |  |  |  |  |  |  |  |
| LC3B | -1.4±0.1 | +15.0 | -4.47 | -5.87 | 0.02±0.01 | 50 | 1* |
| G<sub>ABARAP</sub> | -6.7±0.1 | +1.07 | -0.32 | -6.97 | 0.13±0.01 | 7.75 | 1.17±0.04 |
| PLEKHM1-LIR VNV-CIL |  |  |  |  |  |  |  |
| LC3B | -7.9±0.1 | +3.30 | -0.98 | -8.84 | 3.03±0.27 | 0.33 | 1.06±0.01 |
| G<sub>ABARAP</sub> | -9.2±0.1 | +1.49 | -0.44 | -9.62 | 11.3±0.38 | 0.09 | 1.07±0.01 |

To probe whether PlekHM1 has a preference for the GABARAP family *in vivo* we overexpressed GFP-tagged human ATG8s in the absence and presence of Flag-tagged wild type PLEKHM1 protein (PLEKHM1-WT-Flag) in HEK293T cells. Immunoprecipitation of GFP-mATG8s revealed that PLEKHM1 strongly co-precipitated with LC3C, GABARAP, GABARAP-L1 and GABARAP-L2 and weakly with LC3A and LC3B (Figure 2B). This was recapitulated on an endogenous level in HeLa cells overexpressing GFP-mATG8s and treated with the mTOR inhibitor Ku-0063794 plus chloroquine (Ku+CQ) (Figure 2C), where endogenous PLEKHM1 immunoprecipitated preferentially with GFP-GABARAP and GABARAP-L1 (Figure 2C). Endogenous p62/SQSTM1 co-precipitated with all LC3/GABARAP to a similar extent. Moreover, using either *Plekhm1*^+/+^ or *Plekhm1*^−/−^ (where autophagy is blocked) mouse embryonic fibroblasts, we were able to show that PLEKHM1 and GABARAP, but not LC3B, formed an endogenous complex when PLEKHM1 was immunoprecipitated after Ku+CQ treatment (Figure 2D). This interaction was dependent on PLEKHM1-LIR, as incubation with a Tat-tagged-PLEKHM1-LIR peptide blocked the interaction but not a scrambled control (Figure 2D). Taken together, these data suggest that PLEKHM1 interacts specifically with GABARAP, but not with LC3B, either *in vitro* or *in vivo*, consistent with the hypothesis that its role is as an adaptor and not a receptor protein (Figure 2E).

### Understanding the contributing factors to PLEKHM1-LIR specificity towards GABARAPs

To provide a molecular basis for the specificity of the PLEKHM1-LIR interaction with the mATG8 proteins, we solved the crystal structures of PLEKHM1-LIR in complex with the LC3A, LC3C, GABARAP and GABARAP-L1 proteins. In addition, we included in our comparative analysis the structure of the PLEKHM1-LIR:LC3B complex [PDB: 3X0W (McEwan et al., 2015)]. Thus, we compared the binding of the same LIR motif across multiple members from both the LC3- and GABARAP-subfamilies, an analysis that has not been performed before. To obtain the complex structures, we created chimeric proteins consisting of the mATG8 C-terminally fused to the LIR sequence with a Gly/Ser linker. Crystals diffracted to 2.50 Å for PLEKHM1^629-638^-LC3A^2-121^, 2.00 Å for PLEKHM1^629-638^-GABARAP^2-117^, and 2.90 Å for PLEKHM1^629-638^-GABARAP-L1^2-117^. LC3C could not be crystallized as a chimeric construct, but co-crystals of LC3C with the PLEKHM1-LIR peptide (residues 629-642) diffracted to 2.19 Å resolution. An overview of the structures is provided in **Expanded View Figure EV3 and Expanded View Table EV1**. A detailed analysis of the differences across the LC3/GABARAP proteins is provided in the **Expanded View Results** and summarized below.

We compared the LIR bound and unbound GABARAP family structures to the LC3 family of structures to assess whether global conformational changes account for the preference of PLEKHM1-LIR towards GABARAP. The structures of the PLEKHM1-LIR bound mATG8 proteins overlay very closely (**Expanded View Figure EV4A**), and exhibit conventional pattern of LIR:mATG8 interactions, although subtle differences were observed (**Expanded View Figure EV4C-G** and **Expanded View Table EV2)**.

Next, we analyzed the microenvironment surrounding the four key PLEKHM1-LIR residues W635, V636, N637 and V638 consisting of the core Θ-X_r_X_2_-Γ motif when bound to mATG8 proteins. The HP1 and HP2 pockets are known to be critical for the LIR interaction, and the tighter packing of the two essential residues W635 and V638 into HP1 (Θ) and HP2 (Γ) of GABARAPs versus LC3 families (Figure 3A and 3D and **Expanded View Results**), may in part explain the generally stronger binding of PLEKHM1-LIR to GABARAP proteins.

**Figure 3.**
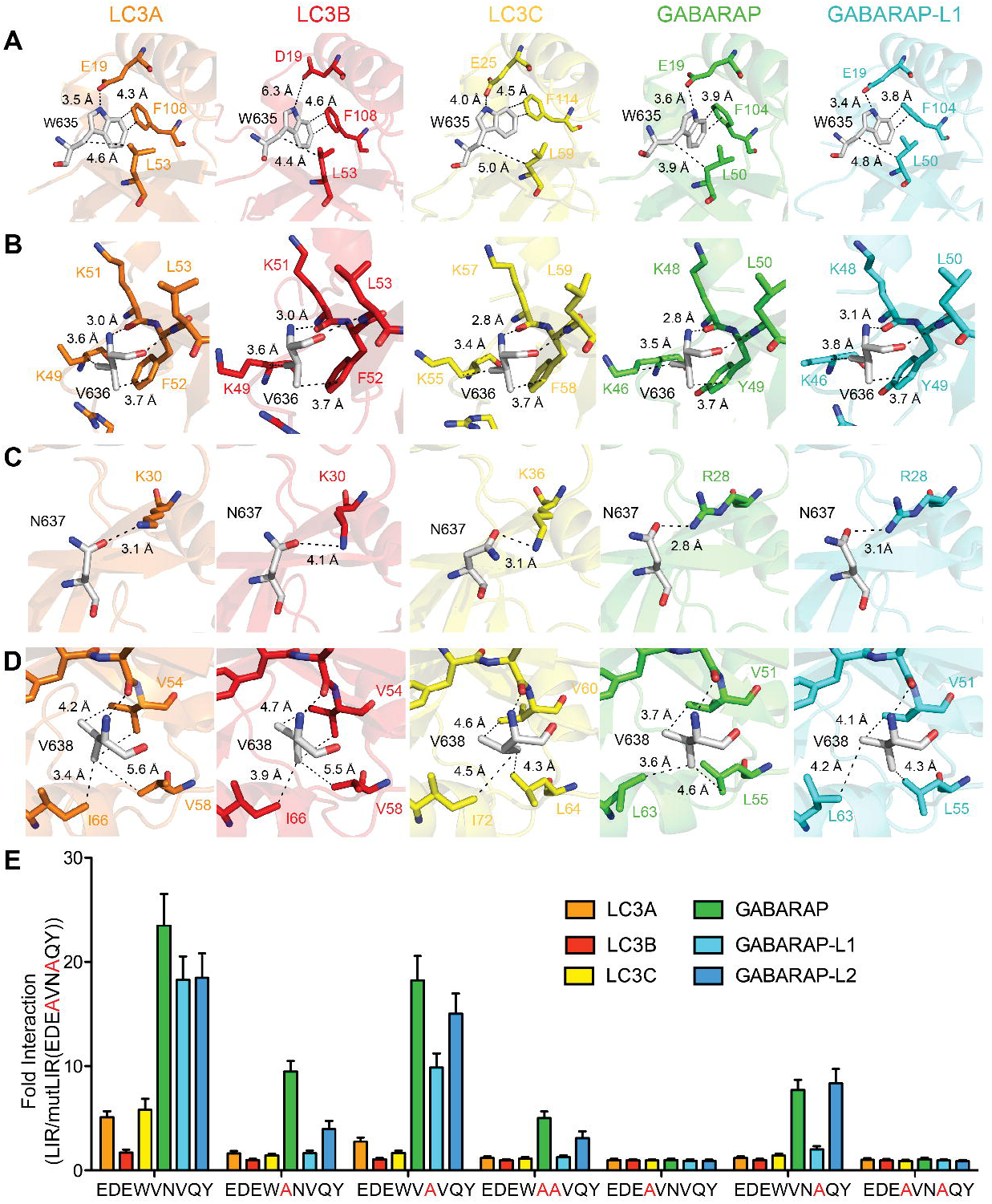
Importance of residues in PLEKHM1-LIR for preferential binding of GABARAP-subfamily proteins. **A.** Sections of the complex structures representing W635 of PLEKHM1 and its microenvironment. Network of intermolecular contacts for the PLEKHM1-LIR residue W635 within complexes with each mATG8 proteins (indicated on each plot). Partner residues in each mATG8 protein are given, hydrogen bounds are shown as dashed lines with distances between corresponding heavy atoms. W635 of PLEKHM1 interacts with E19 of LC3A with 3.5 Å distance, with D19 of LC3B and E25 in LC3C in similar way, but the bond distances are higher (6.3 Å and 4.0 Å), suggesting a weaker interaction. W635 interacts with E17 in GABARAP and GABARAP-L1 with distances of 3.6 Å and 3.4 Å. Additionally, aromatic carbons of W635 are significantly closer to the carbons of GABARAP non-polar residues, forming the HP1. **B.** Sections of complexes structure representing V636 of PLEKHM1 and its microenvironments. V636 in the X_1_ position of PLEKHM1 interacts with residues at the surface of the mATG8 protein. This includes hydrophobic interactions with the aromatic residue (phenylalanine in LC3 and tyrosine in GABARAP) and lysine for both families and for the LC3 protein an arginine also forms part of the interaction surface with V636. In contrast, this arginine in the GABARAP family of proteins is further away and more disordered. **C.** Sections of complexes structure representing N637 of PLEKHM1 and its microenvironments. For the LC3-subfamily proteins, the hydrogen bonding distance of N637 (LIR) with K30 (LC3) correlates with binding affinity to the LIR peptide. The bond distances are (average for all monomers in ASU): 4.2 Å for LC3B, *K*_D_=6.3 μM; 3.1 Å for LC3A, *K*_D_=4.2 μM; and, 3.1 Å for LC3C, *K*_D_=3.5 μM. For LC3A, two of the four monomers in the ASU do not show this interaction and in LC3C, one of the eight monomers in the ASU do not have this interaction, suggesting that this interaction is variable in the LC3 structures. In comparison, R28 in the GABARAP-subfamily proteins is always hydrogen bonded to N637 with a generally shorter bond distance and the geometry of the hydrogen bond between the arginine and asparagine is close to optimal for a hydrogen bond (N-H…O angles are: LC3A - 16.6°, LC3B - 48.3°, LC3C - 26.7°; GABARAP - 4.7°, GABARAP-L1 - 6.9°). The average hydrogen bond distance for all monomers in ASU is: 2.8 Å for GABARAP, *K*_D_=0.6 μM; and 2.8 Å for GABARAP-L1, *K*_D_=0.8 μM. **D.** Sections of complexes structure representing V638 of PLEKHM1 and its microenvironments. Tighter packing of V638 in HP2 of GABARAP-subfamily proteins is observed. V51 GABARAPs sidechain are in close proximity to the PLEKHM1 V638 (3.7 and 4.1 Å for GABARAP and GABARAP-L1, respectively), while side chains of residues in equivalent positions of LC3-subfamily proteins are further away (LC3A/B/C V54/V54/V60 - 4.24.7/4.6 Å). Similarly, GABARAPs L55 sidechain are closer to the PLEKHM1 V638 (4.6 and 4.3 Å for GABARAP and GABARAP-L1, respectively); LC3A/LC3B/LC3C V58/V58/L64 are distanced to PLEKHM1 V638 at 5.6/5.5/4.3 Å. Additionally, the V638 side chain shows some rotational flexibility, observed for when comparing all the crystal structures. **E.** Biotinylated PLEKHM1-LIR peptides (WT and alanine substitutions of highlighted residues) were incubated with streptavidin coated plates, washed and subsequently incubated with 6xHis-tagged mATG8 proteins (human LC3A, -B, -C, GABARAP, -L1 and -L2 proteins). These were washed and incubated with anti-HIS-HRP to detect His-tagged -mATG8s directly bound to biotinylated PLEKHM1-LIR peptides. Samples were again washed and incubated with TMB substrate (3,3’,5,5’-Tetramethylbenzidine). After 5 minutes of incubation time, the reaction was stopped by addition of acid and the samples absorption was directly read at 450nm. Results were normalised to absorbance of the PLEKHM1-mutLIR (EDE<u>**A**</u>VN<u>**A**</u>QY) where both hydrophobic core residues were substituted with alanine and expressed as a fold change of mutant LIR (background noise).

Our structural analysis revealed that PLEKHM1-LIR residues in positions of X_1_ and X_2_ also participate in the binding and could be important for the subfamily-specific interaction’s network (Figure 3B, 3C, **Expanded View Figure EV5** and **Expanded View Results)**. Residue N637 at the X_2_ position forms more preferential contacts for binding of the GABARAP proteins due to better geometry of an intermolecular hydrogen bond to the invariant arginine residue in all GABARAP proteins that is lysine in all LC3 proteins (Figure 3C, **Expanded View Figure EV5B and Expanded View Table EV3**). In contrast, for the V636 in the X_1_ position, we do not observe significant differences in the intermolecular contacts (Figure 3B); however, we observed that in all LC3-subfamily proteins conformation of the V636 is stabilized by intramolecular salt bridge, which is absent in GABARAP-subfamily proteins (**Expanded View Figure EV5A**).

Taken together, our structural analysis reveals that residues in PLEKHM1-LIR positions Θ, Γ and X_2_ form GABARAP-subfamily favorable contacts, while V636 in the X_1_ position has LC3-subfamily favored contacts.

### X_1_ and X_2_ residues are important for PLEKHM1-LIR:mATG8 interaction

To complement our structural studies, we performed a peptide array analysis of the PLEKHM1-LIR mutating each position to analyse the contributing factors of the interactions and how selectivity could be achieved. PLEKHM1-LIR WT peptide (EDEWVNVQY) reproducibly reflected the ITC data (Figure 2A and Table 3) where PLEKHM1-LIR WT with GABARAP (green bar) shows the most potent interaction, followed by GABARAP-L1, -L2, LC3C and LC3A, with LC3B as the weakest interactor (Figure 3E). W635A was sufficient to abolish all PLEKHM1-LIR:mATG8 interactions (Figure 3E), V638A abolished LIR-LC3 family as well as LIR-GABARAP-L1 interactions, but only reduced GABARAP and GABARAP-L2 interactions (Figure 3E), and W635A/V638A completely disrupted all LIR-mATG8 interactions (Figure 3E). Therefore, we are confident that our experimental set-up can be used to accurately assess any alterations in LIR:mATG8 interactions introduced by mutation.

Through substitution of W635 and V638 for residues found in other LIR sequences, we show that W635F and W635Y mutants weaken the interaction with all six mATG8s (**Expanded View Figure EV6A**) but V638L or V638I dramatically increases the binding of PLEKHM1-LIR to LC3B, but did not affect the GABARAP family interactions (**Expanded View Figure EV6A**). Overall, W635 and V638 act as the corner stones for LIR-mATG8 interaction, where W is optimal for all mATG8s and L or I is preferential for LC3B, potentially due to the bulkier residues present in α-helix 3 of GABARAPs that create a shallower HP2 on GABARAPs compared to LC3s, requiring shorter amino acid side chains (V) of the LIR in this position (**Expanded View Figure EV6B-C**).

Next, we assessed the effect of mutation of the X_1_ and X_2_ residues, V636 and N637 respectively, on the interactions of PLEKHM1-LIR with mATG8s. Surprisingly, V636A mutation has a similar effect as V638A, where both significantly reduce the interaction of PLEKHM1-LIR with all LC3 and GABARAP-L1 but reduce GABARAP and GABARAP-L2 interactions (Figure 3E). On the other hand, N637A mutation has a mild effect on GABARAP interactions but strongly reduced LC3A, LC3B and LC3C interactions (Figure 3E).

Taken together, our data indicates that residues in PLEKHM1-LIR positions X_1_ and X_2_ may provide a means of fine tuning the selectivity of LIRs towards LC3 or GABARAP subfamilies.

### Residues at positions X_1_ and X_2_ provide refinement of selective LIR-mATG8 interactions

To study the role of the amino acids in position X_1_ and X_2_ in more depth, we substituted V636 (**Expanded View Figure EV7A and EV7B**) and N637 (**Expanded View Figure EV7C and EV7D**) of PLEKHM1-LIR with all others 19 amino acids and analysed the relative affinity of the each mutated peptide to all six mATG8 proteins in our peptide array (normalizing the strength of interaction in each individual case to that for PLEKHM1-LIR WT). We included the W635A/V638A double mutant (PLEKHM1-mutLIR) as a negative control (Figure 3E). This allowed us to assess mutations that either increased or decreased the interaction with each mATG8 subfamily member, relative to the PLEKHM1-LIR WT sequence. Firstly, we found that substitution of V636 had for most residue types a negative influence on both LC3 (**Expanded View Figure EV7A**) and GABARAP (**Expanded View Figure EV7B**) family interactions, particularly when mutated to G, K, R, P or S (**Expanded View Figure EV7A-B**), indicating that the amino acid in position X_1_ can have a profound impact on LIR-mATG8 interactions. For V636G we confirmed these data by ITC (Figure 4A). Notably, V636C was the only mutant that increased its interaction with any mATG8, specifically LC3B, but did not affect overall interactions with LC3A, LC3C or GABARAP family members (**Expanded View Figure EV7A-B**). Next, we tested the effect of mutating N637 (X_2_) of PLEKHM1-LIR. Overall, substitution of N637 to G or P completely disrupted LIR-LC3 family interactions, with only a mild effect on all GABARAPs (Figure 4A; **Expanded View Figure EV7A-D**). Surprisingly, we found that mutation of N637 to either C, F, I, L, V, W or Y enhanced the interaction of PLEKHM1-LIR with LC3B alone but not LC3A, LC3C or GABARAP family proteins (**Expanded View Figure EV7C-D**).

**Figure 4.**
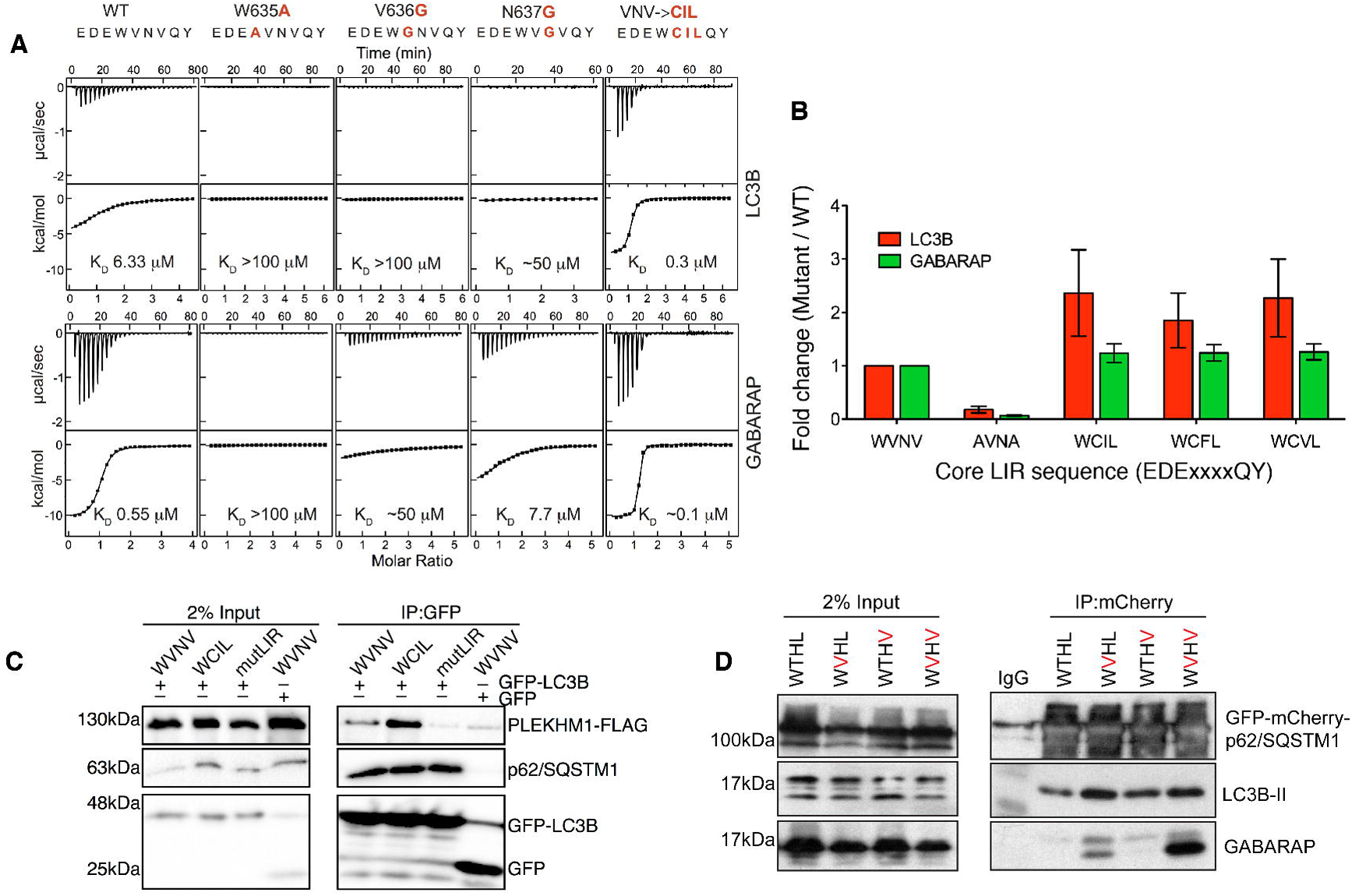
Mutation of X_1_ and X_2_ positions differentially affect interaction with LC3 and GABARAP proteins. **A.** ITC titrations of mutated PLEKHM1-LIR peptide into LC3B (top panel) and GABARAP (bottom panel) proteins. The top diagrams in each ITC plot display the raw measurements and the bottom diagrams show the integrated heat per titration step. Best fit is presented as a solid line. Mutations within the PLEKHM1-LIR peptide are indicated at the top of the figure. **B.** Biotinylated peptides of PLEKHM1- LIR WT (EDE**WVNV**QY), PLEKHM1-mutLIR (EDE**A**VN**A**QY) or mutants that increase LC3B interaction (EDE**WCIL**QY; EDE**WCFL**QY; EDE**WCVL**QY) **C.** Co-immunoprecipitation of GFP-alone or GFP-LC3B with PLEKHM1-WT-Flag, mutant LIR (mutLIR; EDEWVNV/AAAAVNG) or variant LIR (WVNV/WCIL). **D.** GFP-mCherry-p62/SQSTM1 WT, T339V, L341V and T339V/L341V were overexpressed in HEK293 cells and immunoprecipitated using anti-RFP beads and subjected to SDS_PAGE and western blotting. Blots were probed for the presence of endogenous GABARAP and LC3B proteins.

Using combinations of amino acid that individually increased PLEKHM1-LIR:LC3B interaction, we could show that mutation of the core WVNV motif to either WCIL, WCFL or WCVL increased the interaction PLEKHM1-LIR with LC3B (Figure 4B). Indeed, using a WCIL core sequence resulted in a 5-fold increase in GABARAP interaction but a greater than 20-fold increase in the LC3B interaction (K_D_ 0.3μM) (Figure 4A-B). This was mirrored *in vivo* with the PLEKHM1-WCIL (full length) showing increased co-precipitation with GFP-LC3B from cell-lysates compared to PLEKHM1-LIR WT and PLEKHM1-mutLIR (Figure 4C).

Finally, using the established autophagy receptor protein p62/SQSTM1 as a model LIR (DDD<u>**W**TH**L**</u>SS) that interacts with LC3B strongly (K_D_ ~ 1.5μM), we tested whether substitution of T339V (X_1_) and L341V (Γ) altered the selectivity of p62/SQSTM1 LIR *in vivo.* Therefore, we immunoprecipitated GFP-mCherry-tagged wild type and mutant forms of p62/SQSTM1 from HEK293 cells. Under basal conditions, p62/SQSTM1-WT co-precipitated with endogenous LC3B and weakly with endogenous GABARAP (Figure 4D). T339V mutant presented a striking shift in the interaction with endogenous GABARAP over LC3B (Figure 4D) with L341V alone having a mild effect (Figure 4D). However, a double T339V/L341V showed a strongly enhanced shift towards GABARAP with only a moderate increase in endogenous LC3B interaction (Figure 4D). Overall, we have shown that LIR residues at the Γ and X_1_ positions are important for defining GABARAP-selective LIR sequences (GABARAP Interaction Motif; GIM) that are found in a number of endogenous proteins. Moreover, we can alter the selectivity of known autophagy adaptors and receptors by introducing valine residues in the Γ and X_1_ positions to drive towards GABARAP or by mutating X_2_ and Γ-positions to enhance LC3 interaction. In conclusion, the previously unassigned X_1_ and X_2_ positions of a classical Θ-X_1_X_2_-Γ sequence are important regulators of LC3- and GABARAP-subfamily selectivity of LIRs.

## Discussion

The process of building, shaping and “filling” an autophagosome requires a large number of proteins with distinct functions. From E1, E2 and E3-like enzymes to kinases and scaffolds and adaptors that build and transport autophagosomes to their destination. At the core of this process are the small ubiquitin-like modifiers, the ATG8-like proteins, that are conjugated onto the growing autophagosome on both the cytosolic and luminal sides of the nascent autophagosome. The critical positioning of these proteins allows them to recruit both adaptors (present on cytosolic side and that are not degraded in an autophagy dependent manner) and receptors (present on luminal side that are degraded along with the cargo) to the autophagosome [31]. In all cases, the interaction with mATG8 proteins is mediated through a direct interaction between a LIR/AIM motif on the receptor/adaptor and two hydrophobic pockets on the ATG8 proteins. This interaction was first described for the prototypical autophagy receptor protein, p62/SQSTM1, that linked autophagy-mediated protein aggregate degradation with MAP1LC3B conjugated on the autophagosome [7]. Since then, there has been a deluge of both adaptors and receptors identified with conserved LIR motifs that conform to the Θ-X_1_-X_2_-Γ motif. These include autophagy adaptor proteins such as PLEKHM1, ULK1/2, TBC1D5, KBTBD6/7, ALFY and JMY and link the autophagosome to various cellular machineries, such as the autophagosome initiation complex and autophagosome-lysosome fusion machinery [17-19, 24, 25, 32]. Autophagy receptors on the other hand include FAM134B, OPTN, TAX1BP1, NDP52 and p62/SQSTM1 and are linked to the direct removal of a variety of cellular structures and processes from pathogen, protein aggregate, peroxisome and mitochondrial removal, ER turnover and removal of ferritin aggregates (reviewed in [3]).

However, despite the ever increasing number of LC3/GABARAP interaction partners identified and perhaps the over reliance on LC3B as the main marker of autophagosomes, there is now emerging distinct roles of each LC3 and GABARAP subfamily. For example, both LC3 and GABARAP families are essential for autophagy flux [28]; however, LC3s were reported to be involved in phagophore extension and GABARAPs required for autophagosome closure [28]. Moreover, GABARAP can activate ULK1 complex to initiate autophagy, irrespective of its conjugation status [27]. Indeed, this is also reflected in *C.elegans* homologues of GABARAP (LGG-1) and LC3 (LGG-2), where LGG-2 interacts with the Unc51/EPG-1 (ULK1/ATG13) and LGG-2 SQST-1 (p62/SQSTM1) [33]. Overall, there appears to be an evolutionary separation of function of LC3 versus GABARAPs where there may be a preference for GABARAPs conjugated to PE on the cytosolic facing autophagosomal membrane to engage adaptors, and LC3 on the luminal side to recruit receptors and cargo. However, there are some interesting exceptions. For example, OPTN-LIR in its unmodified state, clearly shows preference for GABARAP, however when activated through TBK1-phposphorylation at S177, switches to LC3B indicating a potentially interesting functional change between GABARAP and LC3 families [10, 30]. Also, FYCO1 (LC3A specific adaptor) and NBR1 (GABARAP-L1 specific receptor) are other exceptions that require further exploration[26, 29]. Since the initial identification and characterization of the p62/SQSTM1 LIR, there has been little headway in the identification of LC3- or GABARAP-subfamily selective LIR sequences. Currently, there is only one subfamily specific LIR sequence, CLIR, present in NDP52 and TAX1BP1 [11, 20] that specifically mediates the interaction with LC3C.

For the first time, we provide evidence of a GABARAP-selective LIR motif built around the classical Θ-X_1_X_2_-Γ motif. Using a peptide-based array to test interaction profiles of known LIRs, we found that 14 out of 30 tested had a strong preference for GABARAP versus LC3B. These included ULK1/2 and KBTBD6, which had previously been shown to be GABARAP specific [19, 24], and several that previously had not been identified as GABARAP-selective, including JMY and PLEKHM1. Interestingly, PLEKHM1 showed a strong preference for GABARAP versus LC3 despite the apparent similarity of PLEKHM1-LIR (EDEWVNV) with p62/SQSTM1 LIR (DDDWTHL). Notably, the majority of proteins we identified as more selective towards GABARAP, presented with a valine/isoleucine in the X_1_ position and a valine/isoluecine in the Γ position (64%). Indeed, using mutational analysis of the X_1_ and X_2_ positions of PLEKHM1-LIR, which have previously not been linked to LC3 or GABARAP subfamily interactions, we were able to show that residue X_1_ is important for the LC3 and GABARAP interface. For example, substitution of V636 with small G, A, P, S or positively-charged N, K, R and H residues, are generally disruptive to LIR-mATG8 interactions. We found a more favorable microenvironment of PLEKHM1-LIR X_1_ (V636) for LC3-subfamily structures than for GABARAP-subfamily structures (**Figure EV5A and Expanded View Results**). However, we believe that the observed differences do not provide enough energy to shift the preference of PLEKHM1-LIR towards LC3 proteins and are not reproducible for other LIR:GABARAP structures. For example, the structure of KBTBD6-LIR with core sequence W-V-R-V in complex with GABARAP [19] displays similar microenvironment features of V at X_1_ position as PLEKHM1-LIR V636 in complexes with LC3-proteins. Therefore, the microenvironments of V636 are similar in all PLEKHM1-LIR:mATG8 complexes and, when mutated, results in a universal decrease in interaction with all mATG8s (**Figure EV7A-B**). We also show that substitutions at position X_2_ (N637) are less disruptive; however G and P can also decrease most LIR-mATG8 interactions *in vitro.* When we introduce either K or R in the X_2_ position of PLEKHM1-LIR, thereby making it similar to KBTBD6-LIR (DDFWV<u>**R**</u>VAP) that forms an intermolecular hydrogen bond with GABARAP Y25 [19], we observed a reduced interaction with GABARAPs indicating that although similar in sequence, other factors, such as the F in the X_−1_ position, may also influence selectivity. Perhaps the most surprising results were when we mutated X_2_ (N637) to C, F, I, L, V, W or Y, resulting in a large increase in the interaction with LC3B only, compared to WT PLEKHM1-LIR peptide. Indeed, when we rationally mutate the X_1_, X_2_ and the Γ-positions of PLEKHM1-LIR using combinations that increase LC3B interaction, we can achieve a direct 20-fold increase in the interaction with LC3B using ITC as a measurement.

This alteration is not confined to PLEKHM1, as we show that by introducing a single point mutation in the X_1_ position of p62/SQSTM1-LIR, T339V, we can increase the interaction of p62/SQSTM1 with endogenous GABARAP. In addition, we tested the effect of substitution of a recently identified ALS-FTD p62/SQSTM1 mutation (Γ-position, L341V) that has been associated with poor prognosis [34]. We showed that the L341V mutation alone had little effect on LC3/GABARP specific interaction. However, when we combine T339V and L341V (T329V/L341V) the interaction is dramatically switched towards endogenous GABARAP with little or no effect on the interaction with LC3B interaction. Strikingly, a similar theme can be observed in the LGG-1 and LGG-2 C.elegans homologues, where the presence of V in the Γ-position is more favourable for LGG-2 (GABARAP) and L for LGG-1 (LC3) interactions and introducing V into the X1 position can increase the interaction with LGG-1 [33]. This leads us to propose for the first time a subfamily selective LIR sequence that we have termed a GABARAP Interaction Motif (GIM; [W/F]-[V,I]-X_2_-[V,I]) that is more selective towards GABARAP family of proteins. The identification of a GIM (and its separation from the LIM - LC3 Interaction Motif) will allow more precise and directed autophagy research towards understanding adaptor and/or receptors specific function within the life cycle of an autophagosome and the role of mammalian ATG8 paralogues during autophagosome formation, cargo selection, transport and fusion.

## Materials and Methods

### Cloning plasmid preparation

The genes for the truncated LC3A^2-121^, LC3C^8-125^, GABARAP^2-117^, and GABARAP-L1^2-117^ proteins were cloned into pET30ΔSE vector between the *Bam*HI and *Xho*I sites using previously established protocols [35]. The chimeric constructs of the PLEKHM1-LIR attached to the LC3A, GABARAP and GABARAP-L1 proteins were prepared by inserting the oligonucleotide sequence corresponding to the PLEKHM1-LIR peptide (P^629^QQEDEWVNV^638^) and glycine-serine linker into the *Bam*HI site of the pET30ΔSE vector, placing the PLEKHM1-LIR at the N-terminal of the mature chimeric protein (similar to [30, 35]). For the expression of human LC3 and GABARAP proteins for ITC and NMR experiments, plasmids with appropriate modified Ub-leaders in pET vectors were used [36]. Gene, encoding PLEKHM1-LIR peptide, was ordered as synthetic oligonucleotides (Eurofins Genomics GmbH) and cloned into the pET39_Ub63_ vector [36] by NcoI-BamHI restriction sites. After TEV cleavage, the resulting peptide has the amino-acid sequence GAMG-P^629^QQEDEWVNVQYPD^642^, where the first four residues (GAMG) are the cloning artefact.

### Protein expression and purification

The chimeric constructs were expressed as a His-tag fusion protein in *E. coli* BL21(DE3) cells. The cells were induced with 0.3 mM IPTG at OD_600_ 0.6 for 16 hours at 26 °C. The cell pellets were lysed using mechanical sonication in lysis buffer (20 mM Tris-HCl pH 9.0, 100 mM NaCl, 10 mM Imidazole, supplemented with 0.1% Triton X-100). The proteins were purified using Ni-NTA beads (GE Healthcare) and the His-tag was cleaved using thrombin (Invitrogen) at room temperature for 16 hours. The last step was gel filtration chromatography using a Superdex S200/300 GL column (GE Healthcare). The proteins were concentrated using spin concentrators (Vivaspin). For ITC and NMR studies, the non-labelled and stable isotopes labelled LC3 and GABARAP proteins were obtained based on the protocols described elsewhere [29, 36]. Here, *E. coli* NEB T7 Express culture transformed with corresponding plasmids were grown till OD_600 nm_=1.0 and protein expression was induced with 0.2 mM IPTG. The cultures were incubated at 25 °C for 8-12 hours before cell harvesting. Isolation and purification procedures were similar to the reported in [21, 37]. Before experiments, all proteins and peptides were equilibrated with a buffer containing 50 mM Na_2_HPO_4_, 100 mM NaCl at pH 7.0, and supplied with 5 mM protease inhibitors cocktail.

The protocol for preparation of non-labeled and ^13^C,^15^N-labelled PLEKHM1-LIR peptide was slightly modified to achieve highest yield of the peptide. The 50 ml M9 culture was inoculated with NEB T7 cell transformed with pET39_Ub63- PLEKHM1-LIR plasmid and grown overnight at 37°. The collected cells were re-suspended in 2 L of either LB or M9 media contained 1.5 g ^15^N-labelled NH_4_Cl and 3.0 g of ^13^C-labelled glucose. The cultures were grown at 37° till A(600nm)=0.9 and supplied with 1 mM of IPTG to induce Ub63-PLEKHM1-LIR overexpression (3 hours at 37°). After that cells were harvested by centrifugation, re-suspended in buffer contained 50 mM Tris-HCl pH=7.9, 200 mM NaCl, 5% glycerol, 0.1 mg/ml DNAseA and 4 mM protease inhibitors cocktail. After cell lysis by French press, debris was removed by centrifugation and clear supernatant was applied onto the column contained Ni-NTA sepharose equilibrated with the loading buffer (50 mM Tris-HCl pH=7.9, 250 mM NaCl, 1% glycerol and 20 mM Imidazole). Elution was performed with 400 mM Imidazole in the same buffer. An aliquot of pure Ub63- PLEKHM1-LIR fractions was further purified by gel-filtration on Superdex75 26x60 column for control ITC and analytical size exclusion chromatography experiments, remaining fusion protein was processed with the TEV-protease and PLEKHM1-LIR peptide was purified to 95% purity by reverse Ni-NTA chromatography and followed gel-filtration on Superdex75 26x60 column. Peak maximum of peptide was detected at 97 ml (void volume 115 ml). Pure peptide was concentrated in Amicon concentrators with cut-off 3 kDa (> 95% retention).

### Crystallization and data processing

The PLEKHM1^629-638^-LC3A^2-121^, PLEKHM1^629-638^-GABARAP^2-117^ and PLEKHM1^629-638^-GABARAP-L1^2-117^ chimeric proteins were purified and crystallized as N-terminally LIR fused chimeric proteins. The LC3C^8-125^ protein was co-crystallized with the PLEKHM1-LIR peptide (GAMG-P^629^QQEDEWVNVQYPD^642^). Initial crystallization trial was performed using Hampton Research (Crystal screen, Crystal screen cryo, Index and PEG/Ion) and Molecular dimension (JCSG+, Midas, Morpheus, PACT, Clear Screen Strategy 1 and Clear Screen Strategy 1). In all cases the drops included 400 nL of protein (concentrations listed below) and 400 nL of mother liquor. All crystallization experiments were set up at 4 ^o^C.

For PLEKHM1^629-638^-LC3A^2-121^ (10 mg.ml^−1^) crystals were grown in the JCSGplus screen condition H7 (0.2 M ammonium acetate, 0.1 M Bis Tris, pH 5.5, 25% w/v Polyethylene glycol 3,350). Crystals for PLEKHM1^629-638^-GABARAP^2-117^ (9.1 mg.ml^−1^) were grown in the PEG/ion screen condition F5 (4% v/v Tacsimate pH 8.0, 12% w/v Polyethylene glycol 3,350). Crystals for the PLEKHM1^629-638^-GABARAP-L1^2-117^ protein (7.5 mg.ml^−1^) were formed in the PEG/ion screen condition A6 (20% w/v Polyethylene glycol 3,350, 0.2 M NaCl, 8% MPD pH 7.2). The LC3C^8-125^ protein (9.2 mg.mL^−1^) was mixed with the PLEKHM1 peptide (2.4 mg.mL^−1^) in equal volume and incubated for three hours at 4 ^o^C, prior to setting up the crystallization trays. Crystals were formed in the PEG/ion screen condition D5 (0.2 M Potassium Phosphate monobasic, 20% w/v Polyethylene glycol 3,350). The crystals were frozen in liquid N2 prior to data collection.

X-ray diffraction data was collected on the MX2 micro crystallography beamline at the Australian synchrotron (Melbourne, Australia). The data were integrated using XDS [38] and scaled using Aimless [39]. The PLEKHM1^629-638^-GABARAP^2-117^ and PLEKHM1^629-638^-GABARAP-L1^2-117^ structures were solved by molecular replacement using MOLREP [40] and search models 1GNU and 2R2Q, respectively. Phases for the PLEKHM1-LIR:LC3C co-crystal structure where estimated using PHASER [41] and the search model was 3WAM. The solved structures were refined using PHENIX.REFINE [42] and manual refinement was performed using COOT [43]. The images in the work were generated using PyMOL (The PyMOL Molecular Graphics System, Version 1.5.0.4 Schrödinger, LLC).

### Isothermal titration calorimetry (ITC)

All titration experiments were performed at 25 °C using a VP-ITC microcalorimeter (MicroCal Inc., MA, USA). The ITC-data were analysed with the ITC-Origin 7.0 software with a “one-site” binding model. The peptides at concentrations of 0.4 mM were titrated into 0.020 mM LC3 and GABARAP proteins in 26 steps. The proteins and peptides concentrations were calculated from the UV-absorption at 280 nm by Nanodrop spectrophotometer (Thermo Fisher Scientific, DE, USA).

### Nuclear magnetic resonance spectroscopy

All NMR experiments were performed at 298 K on Bruker Avance spectrometers operating at proton frequencies of 500, 600 and 700 MHz. Titration experiments were performed with a 0.18 mM ^15^N-labelled LC3 and GABARAP protein samples to which the non-labelled PLEKHM1-LIR peptide was added stepwise until 4 times excess to LC3 proteins, or 2 times excess to the GABARAP proteins. Backbone HN resonances for selected mATG8 proteins in complex with the PLEKHM1-LIR peptide were assigned using [^15^N-^1^H]-TROSY versions of 3D HNCACB experiment. For assignment of PLEKHM1-LIR peptide backbone HN resonances in complexes with the LC3 and GABARAP proteins, [^15^N-^1^H]-TROSY versions of 3D HNCACB experiment and hCcconh-TOCSY experiment were used.

### Peptide array

Biotinylated peptides (JPT, Germany) were immobilized on streptavidin coated 96-well plates (#436014; Thermo Scientific) in 100μl PBS containing 0.1% Tween-20 (PT) and 1% BSA (PTB) overnight on a shaker at 8°C. After three washing steps with 200μl PT, 100μl of 1μM HIS6-tagged mATG fusion proteins isolated from *E.coli* in PTB were incubated with the immobilized peptides for 1h at 8°C. After three washing steps with 200μl PT, HIS6-ATG8 bound to peptides was detected after 1h incubation with anti-HIS-HRP antibody (JP-A00612; Genscript; 1:5000 in 100μ PTB) with the help of TMB substrate Reagent Set (BD OptEIA; 75μl). The reaction (blue coloration) was stopped by addition of 60μl 1M H3PO4. Samples were analysed on a Synergy H1 ELISA reader from BioTek at 450 nm.

### Immunoprecipitation

Cells (HEK293T, HeLa and *Plekhm1*^+/+^ and *Plekhm1*^−/−^ mouse embryonic fibroblasts) were lysed in NP-40 lysis buffer (50mM TRIS, pH7.5, 120mM NaCl, 1% NP-40) supplemented with Complete® protease inhibitor (Roche). Lysates were passed through a 27G needle, centrifuged at 21000xg and incubated with either anti-GFP agarose (Chromotek), anti-RFP (Chromotek) or anti-PLEKHM1 (SIGMA, HPA025018) plus Protein-A agarose (Roche), washed 3 times in lysis buffer and subjected to SDS_PAGE and western blot. Anti-GFP (Santa Cruz), anti-FlagM2 (SIGMA), anti-p62 (ENZO), anti-LC3B (clone 5F10 Nanotools), anti-GABARAP (Abcam) and anti-PLEKHM1 (SIGMA) were used to detect co-precipitated proteins.

## Acknowledgements

We thank Natalia Rogova for help with protein and peptide sample preparation. Work of VVR, AK, FL and VD was funded by the Center for Biomolecular Magnetic Resonance (BMRZ, Frankfurt); the German Cancer Consortium (DKTK), the Cluster of Excellence Frankfurt “Macromolecular Complexes”; the LOEWE program of the State of Hesse, Germany; and the SFB 1177 “Molecular and Functional Characterization of Selective Autophagy”. Work of DGM was funded by Wellcome Trust Seed award in Science (202061/Z/16/Z). RCJD, ARC and AL were supported in part by Ministry of Business, Innovation and Employment Contract (contract UOCX1208), the Royal Society of New Zealand Marsden Fund (contracts UOC1013 & UOC1506), and United States Army Research Laboratory and United States Army Research Office (contract/grant W911NF-11-1-0481). RCJD and HS were supported by a Japan Society for the Promotion of Science contract through the Royal Society of New Zealand (FY2012).

## Protein Databank Submission

The atomic coordinates and structure factors (PDB codes 5DPR, 5DPW, 5DPS and 5DPT for complexes of PLEKHM1-LIR with LC3A, LC3C, GABARAP and GABARAP-L1, respectively) have been deposited in the Protein Data Bank, Research Collaboratory for Structural Bioinformatics, Rutgers University, New Brunswick, NJ (http://www.rcsb.org/).

## Author Contributions

VVR prepared all samples and determined the NMR structure of the complex and Kd values by NMR and ITC and worote the manuscript; FL performed NMR experiments; AK prepared mutants and performed ITC and NMR titration experiments; AS performed and analysed peptide experiments; ACR, HS, AL, SW and RCJD performed and analysed x-ray crystallography data. VD, ID, RCJD and DGM wrote the manuscript. DGM designed experiments and co-ordinated the manuscript.

